# Targeted control of pneumolysin production by a mobile genetic element in *Streptococcus pneumoniae*

**DOI:** 10.1101/2020.06.16.154153

**Authors:** Emily J. Stevens, Daniel J. Morse, Dora Bonini, Seána Duggan, Tarcisio Brignoli, Mario Recker, John A. Lees, Nicholas J. Croucher, Stephen Bentley, Daniel J. Wilson, Sarah G. Earle, Robert Dixon, Angela Nobbs, Howard Jenkinson, Tim van Opijnen, Derek Thibault, Oliver J. Wilkinson, Mark S. Dillingham, Simon Carlile, Rachel M. McLoughlin, Ruth C. Massey

**Author notes:** Contributed equally to this work (Emily J. Stevens, Daniel J. Morse, Dora Bonini).

## Abstract

*Streptococcus pneumoniae* is a major human pathogen that can cause severe invasive diseases such as pneumonia, septicaemia and meningitis. Young children are at a particularly high risk, with an estimated half a million deaths worldwide in those under five attributable to invasive pneumococcal disease each year. The cytolytic toxin pneumolysin (Ply) is a primary virulence factor for this bacterium, yet despite its key role in pathogenesis, immune evasion, and transmission, the regulation of Ply production is not well defined. Using a genome-wide association approach we identified a large number of potential affectors of Ply activity, including a gene acquired horizontally on the antibiotic resistance conferring Integrative and Conjugative Element (ICE) ICE*Sp*23FST81. This gene encodes a novel modular protein, ZomB, which has an N-terminal UvrD-like helicase domain followed by two Cas4-like domains with potent ATP-dependent nuclease activity. We found the regulatory effect of ZomB to be specific for the *ply* operon, potentially mediated by its high affinity for the BOX repeats encoded therein. Using a murine model of pneumococcal colonisation, we further demonstrate that a ZomB mutant strain colonises both the upper respiratory tract and lungs at higher levels when compared to the wild type strain. While the antibiotic resistance conferring aspects of ICE*Sp*23FST81 is often credited with contributing to the success of the *S. pneumoniae* lineages that acquire it, its ability to control the expression of a major virulence factor implicated in bacterial transmission is also likely to have played an important role.

## Introduction

For opportunistic pathogens, such as *Streptococcus pneumoniae*, there is a fine balance to be reached between the ability to colonise the host asymptomatically, to transmit between hosts, and to cause disease symptoms [1–3]. The secretion of cytolytic toxins is often key to this. *S. pneumoniae*, for example, produces pneumolysin (Ply), a cytolytic pore-forming toxin that binds to cholesterol in the membranes of host cells, where it inserts into the lipid bilayer forming a transmembrane pore that lyses the host cell [4–7]. Ply also affects the host immune system in a complex manner, with evidence for both pro- and anti-inflammatory activity [8–11]. The data surrounding the role of Ply in nasal colonisation is complex, with early studies suggesting it contributed positively to the colonisation process [12], but more recent work showing its expression to inversely correlate with colonisation duration and directly correlate with shedding of *S. pneumoniae* from the nose for transmission to new hosts [13–15]. Given the importance of Ply to many aspects of the biology of *S. pneumoniae* it represents an attractive target for the development of therapeutic intervention [16].

Despite its importance, what is perhaps surprising about Ply is that so little is known about its regulation compared to the virulence factors of similar pathogens, such as *Staphylococcus aureus*. To begin to address this, here we sought to define the genetic basis of Ply activity by analysing a collection of 165 isolates belonging to the *S. pneumoniae* PMEN1 lineage [17]. This globally successful clonal group (also known as vaccine serotype 23F, multilocus sequence type 81) is of historical importance having made an important contribution to the emergence of penicillin non-susceptibility amongst the pneumococci [18]. It has acquired further antibiotic resistance through the acquisition of the Integrative and Conjugative Element (ICE) ICE*Sp*23FST81 [8], which confers resistance to tetracycline, macrolides and chloramphenicol, features believed to have contributed to the success of this lineage. As such it represents an important and relevant lineage on which to base this study.

## Materials and Methods

### Bacterial strains and growth conditions

Clinical isolates used in this study (listed in supp. table 4) had been previously sequenced [17] and all belonged to the PMEN1 clone of which *S. pneumoniae* ATCC 700669 is the reference strain. Strains were grown for 16-24 hours in 5% CO_2_ at 37°C, on either brain-heart infusion (BHI), Todd Hewitt supplemented with 0.5% (w/v) yeast extract (THY), blood agar plates containing 5% (v/v) defibrinated horse blood, or in BHI/THY broth without blood.

### Construction of deletion mutants in *S. pneumoniae*

Genes were deleted in *S. pneumoniae* using linear PCR products as described previously [19]. In brief, a stitch PCR approach using the primers listed in Table 1 were used to generate a single PCR product consisting of 1kb of DNA to either side if the gene to be deleted, with a gene encoding resistance to erythromycin (*ermAM* - amplified from plasmid pVA838) in the centre. To transform the bacteria with this PCR product, the wild type bacteria were grown overnight in 5ml BHI broth, and 2ml of this culture was used to inoculate a pre-warmed tube of 20ml BHI broth. This was incubated for a further 2 hours, and 0.5ml was added to 9.5ml pre-warmed BHI broth and incubated for 30 minutes. 1ml of this sub-culture was transferred to sterile tubes, 10μl of 10μg/ml competence-stimulating peptide-2 (CSP-2) was added each, and these were incubated for 15 minutes. To 200μl aliquots of this, either the PCR product or sterile water was added, and the mixture incubated for a further 2 hours. This was then added to molten BHI agar (10ml) containing 5% (v/v) defibrinated horse blood, allowed to set in a petri dish and incubated for 2 hours. Plates were then overlaid with a further 10ml molten agar containing 5% (v/v) defibrinated horse blood with erythromycin (2μg/ml) to selected for successfully transformed cells. Plates were incubated for up to five days at 37°C, or until colonies appeared within the agar.

**Table 1:**
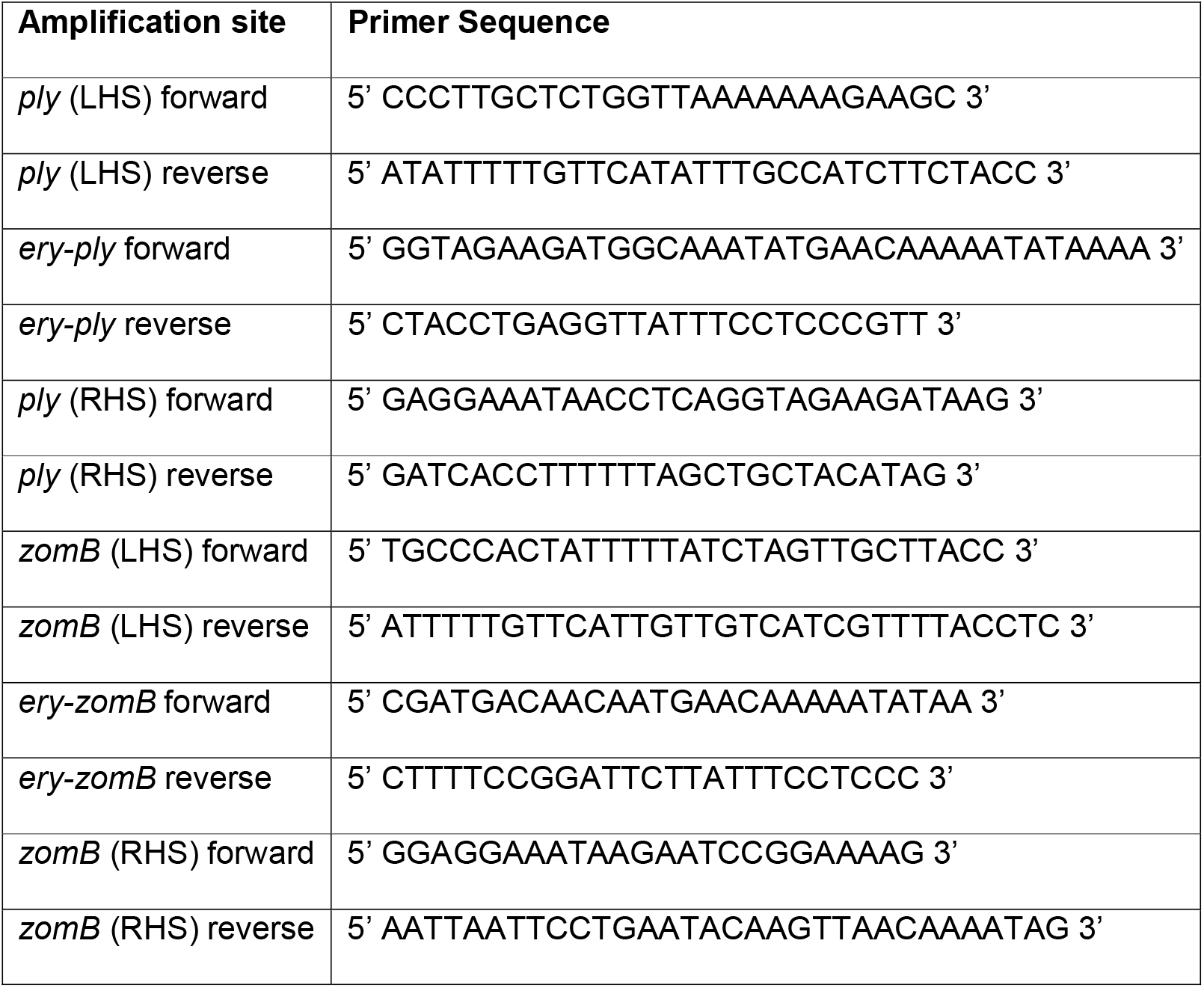
Primers used to construct mutants.

### Cloning of ZomB for complementation

The *zomB* gene was amplified by PCR from ATCC 700669 using KAPA HiFi HotStart ReadyMix (Roche) and primers zomBFW: atatgcatgccctcgtaattactaggaaac (SpHI, Tm 67.6 °C) and zomBRV: atatggatccattttcttatcttatagattctaaaatac (BamHI, Tm 63.7) and cloned into the pVA838 plasmid [20] using MAX Efficiency^™^ DH5α Competent Cells (Invitrogen) to make pVA838-zomB. The plasmid was purified and transformed into D39 as described above, using CSP-1 in place of CSP-2.

### Quantification of Ply activity

Strains were grown overnight in BHI broth, then sub-cultured at a 1:1 ratio into fresh broth and grown for one hour until OD_600nm_ of 0.4-0.7 was reached. These cultures were then serially diluted 4-fold in a 96 well plate containing BSA assay buffer (0.05g bovine serum albumin and 77mg dithiothreitol dissolved in 50ml sterile phosphate buffered saline. To each well, 50μl of triple-washed sheep red blood cells (at a final concentration of 2% diluted in PBS (v/v)) were added and the plates were incubated for one hour in 5% CO2 at 37°C. The plates were then centrifuged at 2000rpm for 10 minutes at room temperature to separate the intact cells from the soluble lysed material. The supernatants from each well were transferred to a fresh 96-well plate and absorbance values read at 415nm were obtained using a FLUOstar Omega microplate reader.

### GWAS

The initial Genome-wide associations between single nucleotide polymorphisms (SNP) and bacterial toxicity were determined by means of linear regression. To account for bacterial population structure, we first performed a singular value decomposition (PCA) of the SNP data and then used the first four principal components (PC), which together explained around 45% of the variance, in the regression model

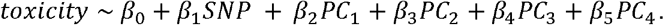

Statistical significance of the *β*_1_ term was determined at an uncorrected *α* = 0.05 threshold. Subsequent GWAS analyses of the data by pyseer and BugWas were performed as previously described [21,22].

### Quantification of Ply production

The bacteria were grown overnight in 15ml BHI broth and harvested by centrifugation. Supernatant proteins were precipitated with 20% trichloroacetic acid, washed 3 times in acetone and resuspended in 100 μl of 8M urea. The individual proteins were separated on an SDS-PAGE gels and Western blotting of this conducted using anti-pneumolysin antibody followed by goat anti-mouse IgG HRP as the secondary antibody. The HRP signal was detected using the Metal Enhanced DAB Substrate Kit (ThermoScientific).

### Quantification of transcription of the *ply* gene

Bacteria were grown overnight in 5-10ml BHI broth and RNA was extracted using a Zymo Quick-RNA Fungal/Bacterial Miniprep kit following the manufacturer’s protocol. Turbo DNase digest kit was used to remove any contaminating DNA. The quality and quantity of the RNA was determined using a Nanodrop and the RNA was reverse transcribed to cDNA using a qScript cDNA synthesis kit (QuantaBio, Beverly, USA), as per the manufacturer’s protocol. RNA samples were standardised such that 100ng was added to each cDNA synthesis reaction. qPCR was performed using a Mic qPCR cycler (Bio Molecular Systems) and reactions were set up using a KAPA Sybr Fast universal master mix (no rox). Three technical repeats were conducted for each cDNA sample and the data analysed using the 2^-(ΔCt ply – ΔCt recA)^ method [23]. The cycle parameters included an initial denaturation at 95°C for 3 min, followed by 35 cycles of 95°C for 10 sec, annealing at 60°C for 20 sec, and elongation at 72°C for 20 sec. A melt curve analysis was performed to check amplified products. The primer pairs used where *recA* was used as the control housekeeping gene:

ply FW: GAAGACCCCAGCAATTCAAG
ply RV: CCTTGAGTTGTTCCATGCTG
recA FW: ATCGGAGATAGCCATGTTGG
recA RV: ATAGAGGCGCCAAGTTTACG

### Expression and purification of ZomB

The *zomB* gene was cloned into the expression plasmid pET15b, and expressed in *E. coli* strain BL21(DE3) with a 6x histidine tag at its N-terminus. The *E. coli* cells were grown at 37°C in LB broth containing 100μg/ml ampicillin to an OD_600nm_ of around 0.5, whereupon cells were temperature acclimatised to 18°C and gene expression was induced for 12 hours via the addition of 1mM IPTG. Cells were harvested by centrifugation for 20 minutes at 6000rpm and resuspended in buffer containing 50mM Tris pH 7.5,150mM NaCl, 10% sucrose and 0.1mM PMSF. Cells were lysed by sonication in buffer containing 1mM TCEP, 500mM NaCl, 20mM imidazole and a protease inhibitor cocktail, and cell debris was removed by centrifugation for 30 minutes at 21,500rpm. Cell lysate was loaded onto a 5ml HisTrap Nickel binding column which had been equilibrated in HisB buffer (20mM Tris pH7.5, 500mM NaCl,1mM TCEP, 5% glycerol) plus 20mM imidazole. Proteins possessing His-tags were eluted with a gradient from 20mM to 500mM imidazole over 32 minutes with a flow rate of 3ml/min. Protein elution was monitored by measuring absorbance at 280nm. Fractions containing the most protein were pooled and applied to a HiTrap Heparin affinity column equilibrated with HepQ/B buffer (20mM TrispH7.5, 1mM TCEP) plus 100mM NaCl. Heparin-binding proteins were eluted with a gradient from 100mM to 1M NaCl over 30 minutes, monitored by absorbance at 280nm. Two millilitre fractions containing the highest protein concentrations were pooled and applied to a MonoQ anion exchange chromatography column equilibrated with HepQ/B buffer plus 100mM NaCl. Proteins were eluted with a gradient from 100mM to 1M NaCl over 30 minutes, with protein absorbance monitored at 280nm, and 0.3ml fractions were collected. Fractions containing the highest concentrations of eluted protein were pooled and stored in HepQ/B buffer plus 330nM NaCl, corresponding to the salt concentration at which the protein was eluted from the column. Nanodrop OD_280_ and the ZomB protein’s predicted extinction coefficient (118390mol/L/cm) were used to determine that ZomB had been purified to a concentration of 7.0μM.

### Biochemical characterisation of ZomB activity

ATPase activity was measured by coupling the hydrolysis of ATP to the oxidation of NADH which gives a change in absorbance at 340nm. Reactions were performed in a buffer containing 20 mM Tris-Cl pH 8.0, 50 mM NaCl, 2 mM DTT, 1 mM MgCl_2_, 50 U/mL lactate dehydrogenase, 50 U/mL pyruvate dehydrogenase, 1 mM PEP and 100 μg/mL NADH. Rates of ATP hydrolysis were measured over 1 min at 25°C and the ssDNA substrate used was Poly(dT). For calculation of K_DNA_ (defined as the concentration of DNA at which ATP hydrolysis is half-maximal), the ATP concentration was fixed at 2 mM. The Michaelis-Menten plot was performed at saturating DNA concentration which is defined as 10x the K_DNA_ value. The concentration of ZomB was 50 nM in these assays unless indicated otherwise.

### Electrophoretic mobility shift assays

The intergenic region between *ply* and the neighbouring gene SPN23F19460, containing BOX repeat regions was amplified (using the following primers; BoxF: GAGAGGAGAATGCTTGCGAC and BoxR: TAGGAATCTCCTTTTTTCACATTTTAATCTTTC). A region of the *ply* gene of equivalent size was also amplified (using plyF: ATGGCAAATAAAGCAGTAAATGACTTTATAC and plyR: GCCCCCTAAAATAACCGCCTTC). The purified ZomB protein was added in a range of concentrations (5μM, 2.5μM, 1.25μM, 0.625μM, 0.3125μM and 0.15625μM) to 10nM of the PCR products in a buffer containing 20mM Tris (pH 8), 200mM sodium chloride, 1mM Tris(2-carboxyethyl) phosphine (TCEP) and 10% glycerol. Samples were then run on a 1.5% agarose gel in 1X TAE buffer for 110 minutes at 90V. Following this, the gel was stained in TAE containing 1X SYBR Safe DNA gel stain for 30 minutes, and bands were visualised using a Typhoon FLA 9500.

### RNA sequencing

Three independent 20 ml cultures of *S. pneumoniae* ATCC 700669 wild type and Δ*zomB* mutant were grown overnight in Todd Hewitt broth supplemented with 0.5% yeast extract, from which total RNA was extracted and DNase treated as described above. RNA was stored at −70°C until transportation on ice for sequencing at the University of Bristol Genomics Facility. RNA integrity was determined by electrophoresis using TapeStation (Agilent) RNA Screentape Assay and samples with scores of > 7 were considered suitable for library preparation and sequencing.

One hundred nanograms of total RNA was taken into the Illumina TruSeq Stranded Total RNA with Illumina Ribo-Zero Plus rRNA Depletion kits according to the manufacturer’s instructions. Briefly, the protocol involved enzymatic depletion of ribosomal RNA and clean-up of the remaining RNA using magnetic beads. The RNA was fragmented and denatured, and first and second strand cDNA was synthesised, then total cDNA purified using magnetic beads. The 3’ ends were adenylated to prevent blunt end ligation, and indexing adapters were ligated to the ends of the double stranded cDNA fragments. Magnetic beads (Agencourt AMPure XP beads, Beckman Coulter) were then used to clean up the cDNA libraries before amplification of DNA fragments, selecting for adapter molecules. A second clean up with magnetic beads was performed and the libraries were quantified using the ThermoFishrer High Sensitivity dsDNA Qubit assay and validated using the TapeStation (Agilent) with the DNA100 screentape assay. The cDNA libraries were normalised to 4nM and pooled for sequencing on the Illumina NextSeq500 instrument using a High Output Version 2.5 sequencing kit. The indices for each sample are detailed in Table 2. The depth of sequencing covered 2 x 75bp paired end reads, with a minimum of 30 million reads per sample. Sequence information was output into FastQ file format for subsequent downstream analysis.

**Table 2:**
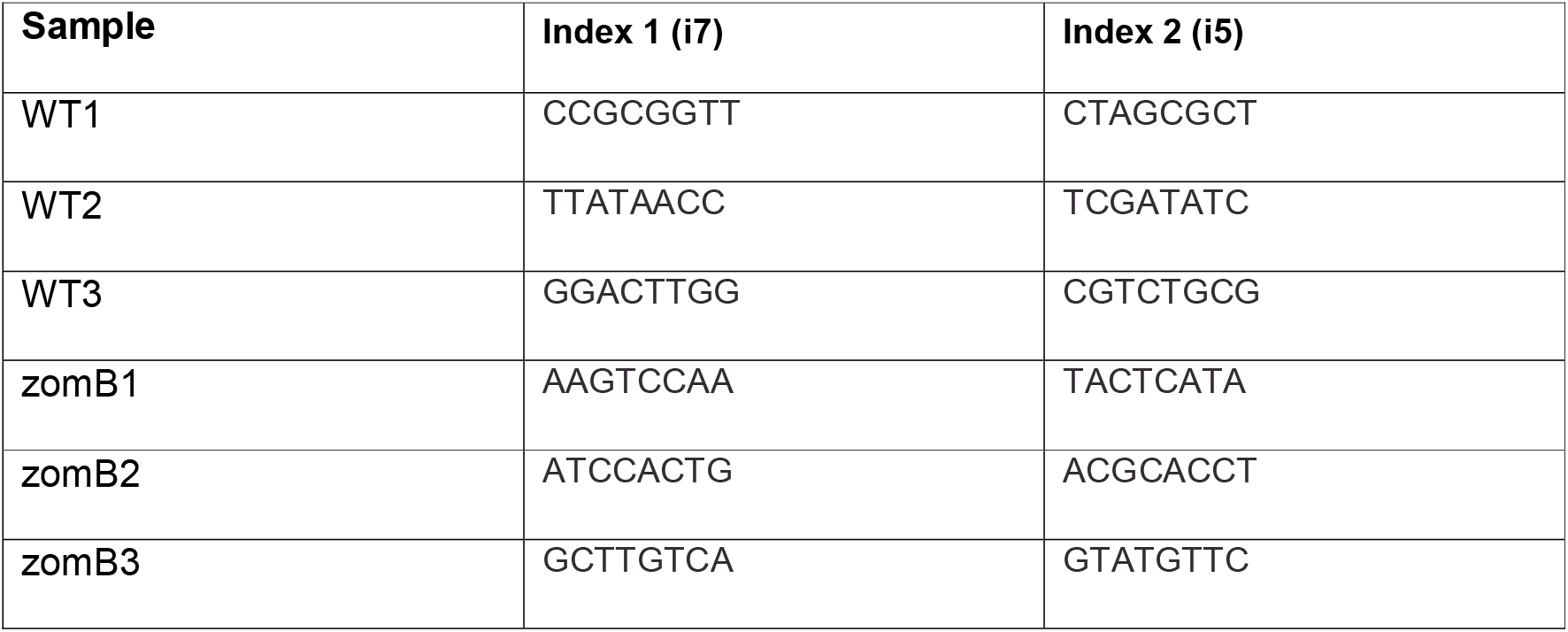
Sample indices for sequencing.

### Bioinformatic analysis of RNA-sequencing data

FASTQ output files from the Illumina sequencing platform (four files for each forward and reverse read (one for each lane used in sequencing)) were concatenated into a single file for each forward and reverse for each sample. The FASTQ files were checked for quality using FASTQC (version 0.11.9, Brabraham Bioinformatics, UK), ensuring consistency and high-quality scores, particularly within ‘per base sequence quality’ and ‘per sequence quality score’ tabs. These files were then taken through a bioinformatics pipeline (via terminal on an Apple MacBook pro running macOS Catalina (version 10.15.7)) detailed below.

An index for the *S. pneumoniae* reference genome (GenBank accession GCA_000026665.1) was created using Bowtie 2 [24] (version 2.4.1), and FASTQ files were aligned to this also using Bowtie 2. Overall alignment rate for the samples was between 87.54% to 97.28%. The annotated genome was then used to create a sequence alignment map (SAM) using SAMtools [25] (version 1.11). Here, a quality control step involved viewing the header of the files to check for quality scores and correct file format. The SAM files were sorted and converted to binary alignment map (BAM) files using SAMtools. The annotated genome was then used as a reference for counting the sorted BAM file reads using Subread featureCounts [26] (version 2.0.1) including options for paired-end reads.

The featureCounts output file was imported into RStudio (Version 1.2.5033) for differential expression analysis using DESeq2 [27] (version 1.26.0). The RStudio source code is available upon request. The aligned BAM files and reference can be obtained at the NCBI Sequence Read Archive (SRA) under accession PRJNA706751.

### Mouse intranasal challenge

8–10-week-old female C57/Black6 mice were inoculated intra-nasally with 1×10^6^ CFU of WT, or an isogenic ZomB deficient strain under isoflurane anaesthetic. Mice were culled at specific time points post inoculation. The upper respiratory tract was lavaged by the insertion of a 20-gauge IV catheter into the trachea with 1ml of sterile PBS washed through and collected at the nose. Lungs were removed and homogenised in 1ml of sterile PBS. Lung homogenate and nasal lavage, where plated on BHI agar with 5% (v/v) defibrinated horse blood and 2.5μg/ml tetracycline for enumeration of CFU.

### Ethics statement

C57/Bl6 mice were bred in-house in Trinity College Dublin. All mice were housed under specific pathogen-free conditions at the Trinity College Dublin Comparative Medicines unit. All mice were used at 8–10 weeks. All animal experiments were conducted in accordance with the recommendations and guidelines of the health product regulatory authority (HPRA), the competent authority in Ireland and in accordance with protocols approved by Trinity College Dublin Animal Research Ethics Committee.

## Results

To examine what variability exists in the production of Ply across a collection of closely related *S. pneumococcal* clinical isolates we first constructed a *ply* mutant in the pneumococcal PMEN1 isolate ATCC 700669 [8] by replacing the *ply* gene with an erythromycin resistance cassette. Using this wild type and mutant strain as positive and negative controls we quantified the Ply activity (lysis of sheep red blood cells (RBCs)) of 165 PMEN1 clinical isolates in triplicate, demonstrating that it varied significantly across this collection of closely related isolates (Fig. 1a). As the genomes of each of these isolates have been sequenced [17], we applied three complementary genome-wide association (GWAS) approaches to identify loci associated with Ply activity. In addition to a linear regression approach using the SNP (single nucleotide polymorphism) data, we also applied two methods that make use of kmer (lengths of nucleotide sequences) data: *BugWAS* [21] and *pyseer* [22]. The results from each of these methods are presented in supplementary tables 1-3. A Manhattan plot in Fig. 1b shows the significantly associated genetic loci determined through the SNP-based method, highlighting those loci where two or all three GWAS methods agreed. (Note, the *P* values are not comparable between the three methods, so this graph is provided as an illustration of the genomic location of the commonly associated loci). There were two notable observations from these analyses. The first was that five loci were associated with Ply activity across all three methods, with the *pbpX* gene, which encodes the Penicillin Binding Protein 2x, being the most significant. This was followed in order of significance by the intergenic region between a gene with the locus tag SPN23F05820 and *bgaA*, a gene with the locus tag SPN23F00840, the intergenic region between a gene with the locus tag SPN23F19120 and *msmG*, and the intergenic region between a gene with the locus tag SPN23F14800 and *greA*. The second notable observation was that 48 individual genes or intergenic regions on the Integrative and Conjugative Element (ICE) ICESp23FST81 were associated with Ply activity.

**Fig. 1:**
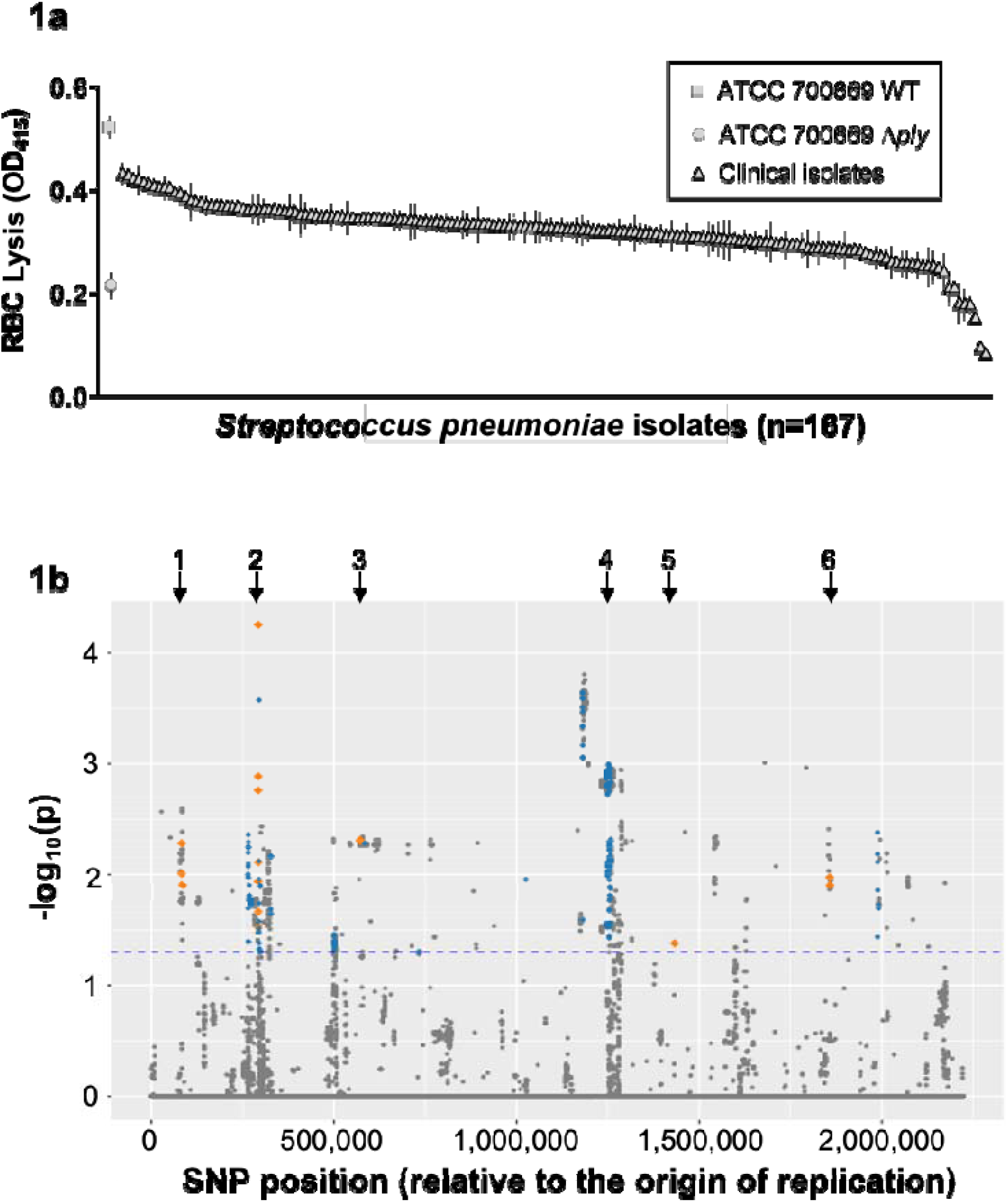
Genome wide association study (GWAS) to identify novel effectors of pneumolysin (Ply) production by *S. pneumoniae*. (**a**) Pneumolysin activity of 165 *S. pneumoniae* of the PMEN1 lineage, as measured by cell lysis. A wild type (WT) and isogenic pneumolysin mutant (Δ*ply*) have been included as controls. (**b**) Manhattan plot of the SNP-based GWAS. The horizontal blue dotted line indicates the threshold for significance (not corrected for multiple tests). SNPs in loci identified by two or all three GWAS methods are indicated in blue and orange respectively. The black arrows indicate the regions of interest; 1: SPN23F00840, 2: *pbpX*, 3: intergenic between SPN23F05820 and *bgaA*, 4: ICESp23FST81, 5: intergenic between SPN23F14800 and *greA*, and 6: intergenic between SPN23F19120 and *msmG*.

Fitness trade-offs between antibiotic resistance and virulence in bacteria are well established [28–31], and for both clinical isolates and isogenic mutants the acquisition of penicillin resistance has been shown to decrease the virulence of *S. pneumoniae* in murine models of infection [32,33]. Given the association of the *pbpX* gene and Ply activity, we hypothesised that a similar trade-off may be occurring here, such that the polymorphisms in the *pbpX* gene may be increasing the levels of resistance to penicillin, which may consequently reduce the levels of Ply being produced, or vice versa. Although bacterial GWAS results are typically validated through mutation of the associated locus, the contribution of the protein encoded by *pbpX* to the biosynthesis of the peptidoglycan layers in the bacterial cell wall is such that it is essential and cannot be inactivated. Instead, we tested our hypothesis by examining the levels of resistance to penicillin (minimum inhibitory concentrations, MICs) for the isolates, but found no significant correlation between the MICs and Ply activity (Pearson product-moment correlation r^2^ = 0.076, P = 0.36). An alternative hypothesis is that the polymorphisms in the *pbpX* gene affects the stem peptide composition of peptidoglycan, as this has been shown to inhibit the release of Ply from the bacterial cells [34]. Further work to test this hypothesis is currently underway.

The high number of associated loci on ICE*Sp*23FST81 is particularly intriguing. This mobile genetic element (MGE), which can be found both integrated and in plasmid form (illustrated in Fig. 2a), is believed to be critical to the success of this lineage of *S. pneumoniae* due to the antibiotic resistance capabilities it brings to the bacteria [8]. Given its ability to move horizontally as a single contiguous unit between bacteria, it is likely that of the associated loci only one is an affector of Ply activity, whilst the others are associated through their physical linkage to this. A survey of the putative activity of all 48 associated loci revealed that the majority of these are genes typical to such elements, involved in antibiotic resistance and the mechanics of its movement. An interesting exception is a gene with the locus tag SPN23F12470. This locus has been annotated as encoding a UvrD-like helicase, a family of proteins typically associated with core house-keeping activities for bacteria. Further *in silico* analysis of the encoded protein, that we have named ZomB, suggests that it is a multi-domain protein with a putative DNA binding helicase domain, followed by two Cas4-like nuclease domains which are predicted to harbour 4Fe-4S clusters [35] (Fig. 2b). With several known examples of Cas-like proteins regulating bacterial virulence [36–38], we hypothesised that it is this gene on ICE*Sp*23FST81 that is the affector of Ply activity.

**Fig. 2:**
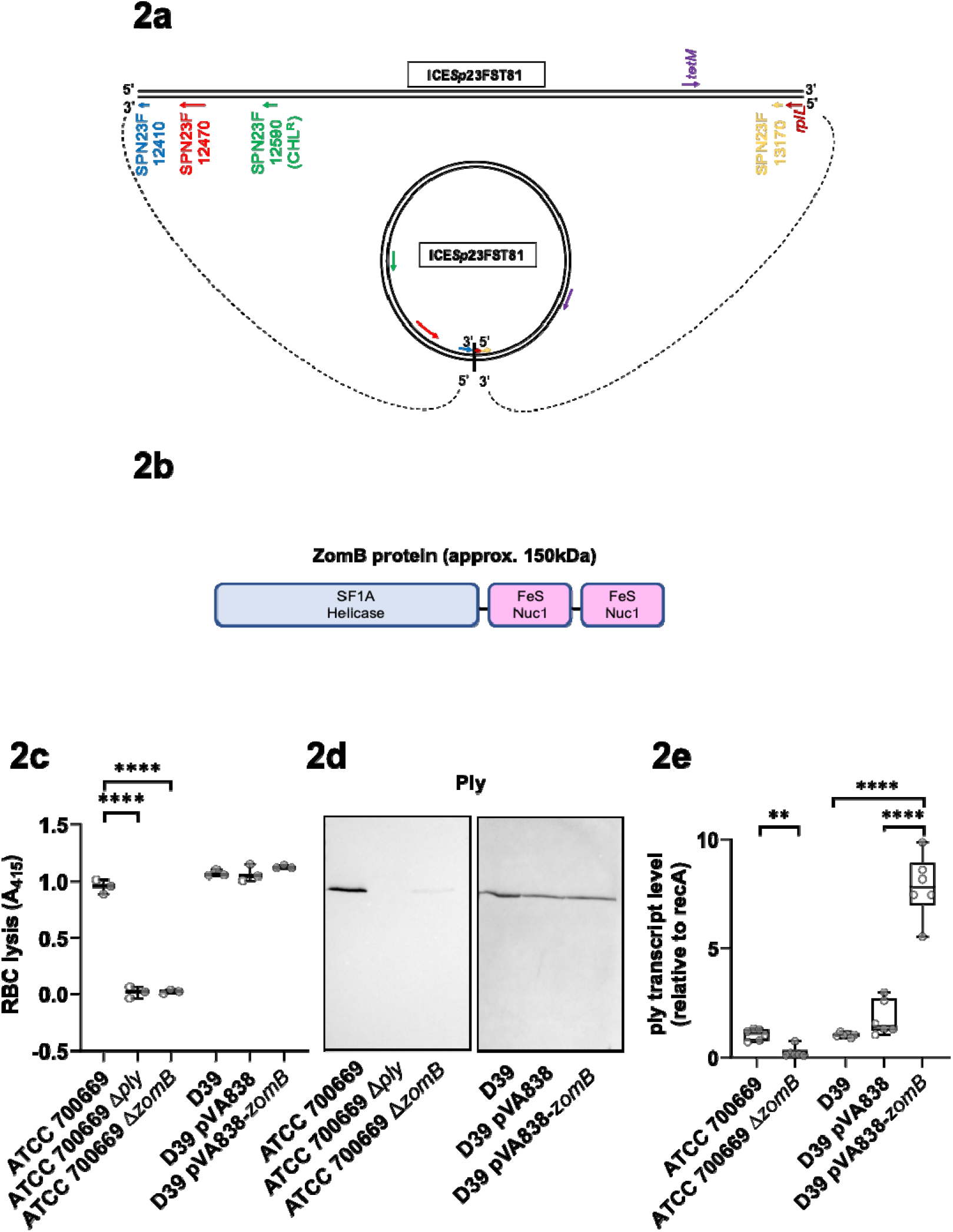
Inactivation of the *zomB* gene on the ICESp23FST81 affects pneumolysin production. **(a)** Cartoon illustration of ICESp23FST81 in both its linear chromosomally integrated form and circularised plasmid form. Some genes of interest have been included: SPN23F12410, SPN23F13170 and *rplL* because these flank the element in its linear form and come into close association when in the plasmid form. Genes encoding the antibiotic resistance genes for chloramphenicol and tetracycline are also indicated. **(b)** Schematic of the ZomB protein with its helicase and two Cas4-like nuclease domains indicated. (**c**) The lytic activity of the *zomB* mutant is comparable to that of the *ply* mutant as determined by a sheep RBC lysis assay in the ATCC 700669 strain. In the D39 strain the introduction of the *zomB* gene on the pVA838 plasmid did not affect the RBC lytic activity of the bacteria. (**d**) The presence of Ply in the extracellular medium is reduced in the *zomB* mutant in the ATCC 700669 strain, determined using anti-Ply antibodies in a Western blot on concentrated bacterial supernatant. In the D39 strain the introduction of the *zomB* gene on the pVA838 plasmid did not affect the level of Ply production. (**e**) The inactivation of *zomB* in the ATCC 700669 strain reduces the transcription of the ply gene, and the introduction of the *zomB* gene into the D39 strain increase *ply* transcription. The transcription of *ply* was determined by qRT-PCR, where the data was made relative to the expression of the housekeeping gene *recA* in each sample and normalised to the level of expression on the *ply* gene in the wild type strain. The box plots represent the median and interquartile ranges; individual data points are indicated by open circles.

To establish the role of ZomB in Ply activity we replaced the *zomB* gene with an erythromycin resistance cassette in *S. pneumoniae* strain ATCC 700669. While no differences in growth between the wild type and mutant strains was observed, the RBC lytic activity of the *zomB* mutant was significantly impaired relative to the wild type strain (Fig. 2c). Using anti-Ply antibodies in a Western blot to understand the mechanism by which ZomB affects Ply activity, we found that this reduction in lytic activity was due to a decrease in the amount of Ply protein being released into the bacterial supernatant (Fig. 2d). We also quantified the relative transcription of the *ply* gene by qRT-PCR and found that the reduced abundance of Ply was due to a significant decrease (19-fold) in *ply* transcription when the *zomB* gene was inactivated (Fig. 2e). Despite numerous attempts to complement this mutation we were unable re-transform the *zomB* mutant with a plasmid containing the *zomB* gene, or the empty pVA838 plasmid. So instead, we introduced it into a more genetically amenable *S. pneumoniae* strain, D39, which does not contain ICE*Sp*23FST81 or the *zomB* gene. The introduction of the *zomB* expressing plasmid to this strain did not increase the ability of the strain to lyse RBCs, or increase its production of Ply (Fig. 2c and 2d). However, the introduction of the *zomB* plasmid significantly increased the transcription of the *ply* gene in D39, verifying the positive effect ZomB has on *ply* expression (Fig. 2e). It is possible that the increase in *ply* transcription did not affect the level of Ply production in this strain due to it being already quite a high Ply producer where it’s protein translational machinery or it’s secretory mechanism may already working at a maximal level.

To characterise the biochemical activities of the ZomB protein, we expressed and purified recombinant ZomB with an N-terminal 6x histidine tag (Fig. 3a). Using a coupled-assay, we found that ZomB hydrolyses ATP with Michaelis-Menten kinetics and displays a turnover number of approximately 20 s^-1^ and a K_m_ value of 90 μM ATP (Fig. 3b). These experiments were performed in the presence of saturating quantities of ssDNA, which was shown to strongly stimulate ATP hydrolysis with an apparent dissociation constant of approximately 1 μM (ntds) (Fig. 3c). This behaviour is typical of the UvrD-like DNA helicases of which ZomB is a member [39]. To examine the putative nuclease activity, ZomB protein (50nM) was incubated with duplex DNA with and without ATP and Mg^2+^. Degradation of the DNA by ZomB was monitored by gel electrophoresis, which showed that the protein possesses a potent ATP-dependent nuclease activity (Fig. 3d).

**Fig. 3:**
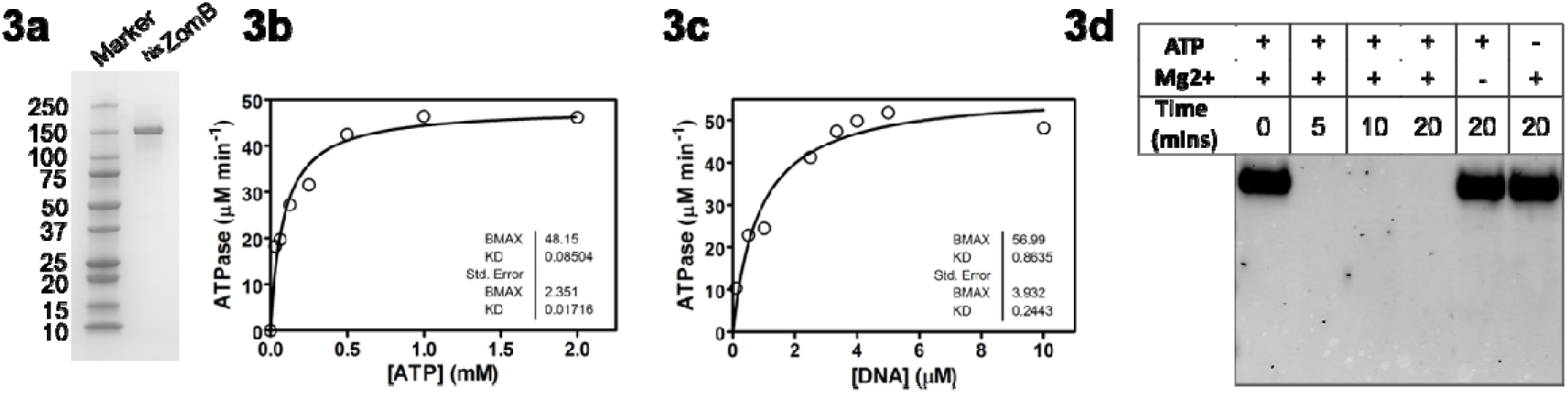
The ZomB protein has ATP dependent nuclease activity. (**a**) An SDS-PAGE gel showing the purified his tagged ZomB protein. (**b**) Steady-state ATPase activity of ZomB (50 nM) was measured at saturating ssDNA concentration to determine the Michaelis-Menten parameters. (**c**) Steady-state ATPase activity of ZomB is strongly stimulated by ssDNA with an apparent dissociation constant of approximately 1 μM ntds. (**d**) Nuclease assays were performed with linear DNA demonstrating that it was degraded by the ZomB protein in the presence of both ATP and divalent cations.

Given the *in vitro* DNA binding and nuclease activity of ZomB, and its role in the transcription of the *ply* gene, we sought to determine the scale of its regulatory activity. To examine this, we compared the level of transcription of all *S. pneumoniae* strain ATCC 700669 coding regions between the wild type and *zomB* mutant using RNA-seq. Under the growth conditions used (i.e. overnight cultures grown in Todd Hewitt broth supplemented with 0.5% yeast extract), we used a >2-fold difference in expression and a P value of <0.05 following Benjamini-Hochberg adjustment as our significance threshold. We found the transcription of nine genes to be affected by the loss of the *zomB* gene, with those genes encoded within the *ply* locus being the most significantly affected (Table 3, Fig. 4).

**Fig. 4:**
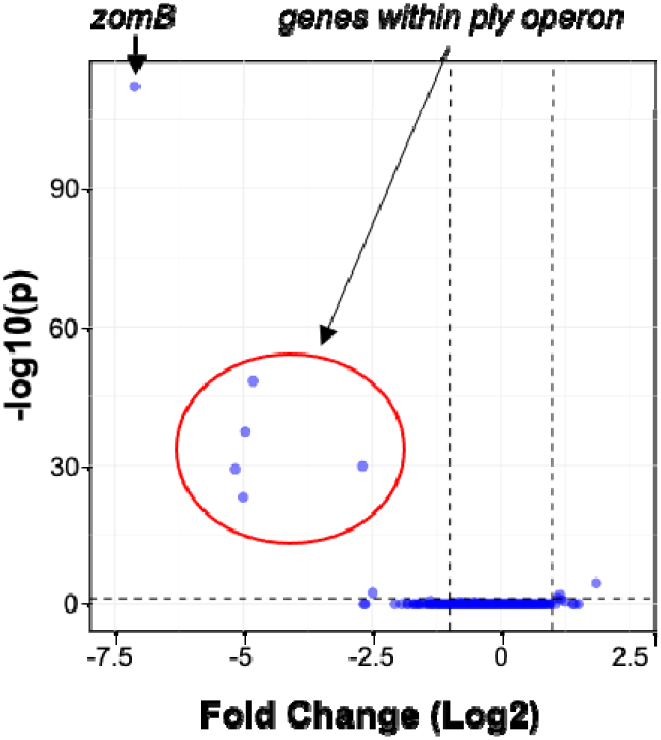
ZomB is a positive regulator of the *ply* operon. The transcription of coding regions across the wild type *S. pneumoniae* and ZomB mutant were compared by RNAseq. Only nine genes were significantly affected, and of those the most affected were *zomB* and the five genes encoded on the *ply* operon.

**Table 3:** Transcriptional differences between the *zomB* mutant relative to the wild type strain. The locus tags in bold indicate those encoded within the *ply* operon.

| Locus Tag | Fold Change (Log2) | P values (adjusted) | Gene Product |
| --- | --- | --- | --- |
| SPN23F11770 | 1.8 | $2.3 \times 10^{-5}$ | ABC-F family ATP-binding cassette domain-containing protein |
| SPN23F12440 | 1.1 | $9.3 \times 10^{-3}$ | Plasmid mobilization relaxosome protein MobC |
| SPN23F12470 | -7.1 | $7.8 \times 10^{-113}$ | ZomB protein |
| **SPN23F19450** | -2.5 | $3.1 \times 10^{-3}$ | MarR family transcriptional regulator |
| **SPN23F19460** | -2.7 | $1.3 \times 10^{-30}$ | YebC DNA-binding transcriptional regulator |
| **SPN23F19470** | -4.8 | $4.4 \times 10^{-49}$ | Pneumolysin |
| **SPN23F19480** | -4.9 | $4.1 \times 10^{-38}$ | Hypothetical protein |
| **SPN23F19490** | -5.2 | $5.5 \times 10^{-30}$ | Hypothetical protein |
| **SPN23F19500** | -5.0 | $6.4 \times 10^{-24}$ | DUF4231 domain-containing protein |

The *ply* gene is transcribed as part of an operon with four other genes that encode a transcriptional regulator (YebC) and proteins implicated in the movement of Ply from the cytosol to the bacterial cell wall (SPN23F19480-19500) (Fig. 5a) [40, 41]. The operon also contains a BOX repeat region immediately downstream of the *ply* gene. Due to their internal repeating sequences, BOX regions can form stable secondary structures, and their presence has been associated with altered transcription of neighbouring genes [42,43]. The molecular details of how their presence affects gene transcription has not yet been determined but is likely due to these secondary structures where they can either enhance or interfere with transcription processes depending on their relative positioning [43]. As ZomB appears to be a positive effector of *ply* transcription, and given its likely DNA binding capability, we hypothesised that it may directly interact with the *ply* locus, perhaps via its BOX region. To test this we amplified two regions of DNA from within the *ply* locus, one containing the BOX element from within the *ply* operon, as well as an equivalently-sized region of DNA within the *ply* coding region, and performed electrophoretic mobility shift assays (EMSAs) with increasing concentrations of ZomB protein. As visualised in Fig. 5b & 5c, the ZomB protein caused a shift in size of both regions of DNA with a higher level of affinity for the BOX containing DNA evidenced by the shift occurring at lower concentrations of protein and with a clearer (less fuzzy) shift in the DNA. While it is tempting to speculate from this that the effect of ZomB on the transcription of the *ply* locus might be mediated via the BOX region, that the transcription of none of the genes neighbouring the other 136 BOX regions scattered across the *S. pneumoniae* genome were affected by the loss of the *zomB* gene does not support this (Table 1 Fig. 4). But what is clear from this analysis is that the ZomB protein can bind to at least two regions within the *ply* locus with a high affinity, and this is likely to be the means by which it elicits its effect on the transcription of these genes.

**Fig. 5:**
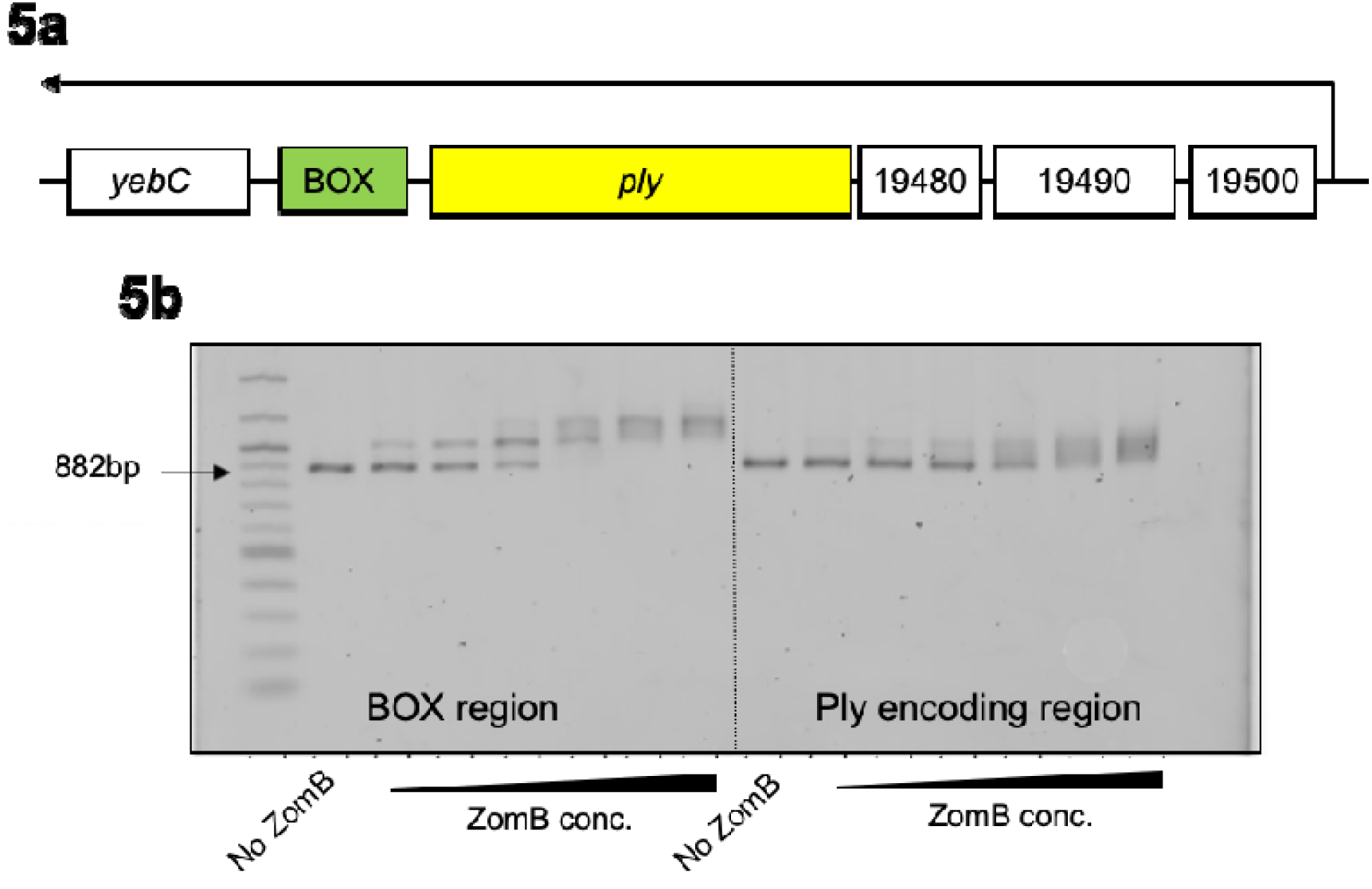
The ZomB protein specifically binds to the BOX region within the Ply encoding operon. (**a**) cartoon of the *ply* encoding operon with the *yebC* and *ply* genes, the SPN23F-locus tags of the neighbouring genes, and promoter and transcript length (black arrow) indicated. (**b**) EMSAs demonstrating the specificity of binding of the ZomB protein for two regions within the *ply* locus. The concentrations of ZomB used were 2.5, 5, 10, 20, 40 and 80nM.

The ability of pathogens to alter expression of key virulence factors to counteract host immune responses is a critical strategy of disease tolerance employed by obligate symbionts, such as *S. pneumoniae*, to facilitate persistence within its host [44,45]. In the absence of Ply a reduced local pro-inflammatory response [13,46] coupled with the potential for a more intracellular lifestyle [41,47] has been shown to promote persistence both in the nasopharynx and in the lungs. With Ply expression inversely correlated with colonisation, we sought to determine how ZomB, a regulator of *ply* transcription, would affect *S. pneumoniae* colonisation in a murine model. Mice were inoculated intra-nasally with a sub-lethal dose of either the wild type or ZomB mutant and nasopharyngeal bacterial burden monitored over 7 days (Fig. 6). The ZomB mutant demonstrated increased persistence within the nasopharynx compared to the wild type strain with increased numbers of ZomB mutant bacteria recovered from the upper respiratory tract at days 3 and 7 post inoculation (Fig 6a). Consistent with this, significantly increased levels of pneumococci were also recovered from the lungs of the ZomB mutant challenged animals compared to animals challenged with the wild-type strains on day 7 post colonisation (Fig 6b).

**Fig. 6:**
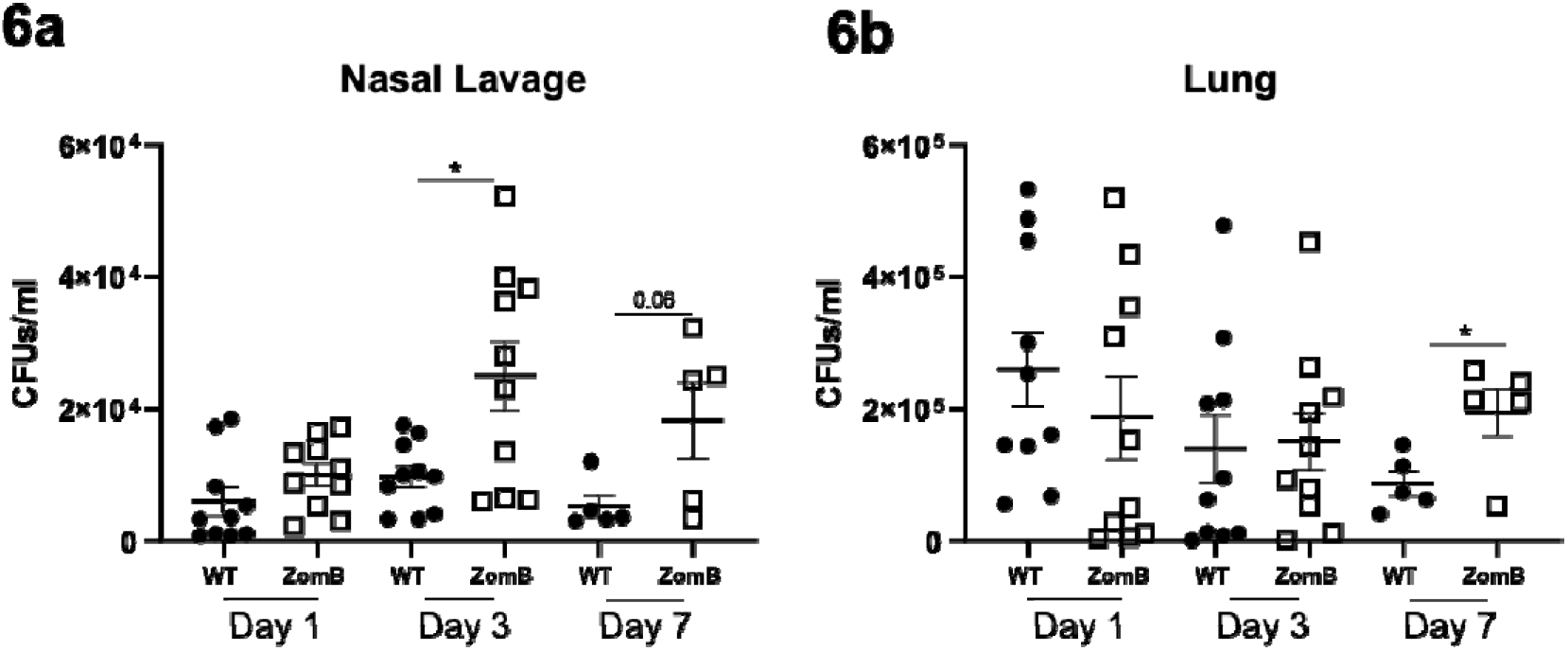
ZomB has a negative effect on the nasal colonisation of mice by *S. pneumoniae*. (**a**) Groups of C57Blk mice were inoculated intranasally with wild type *S. pneumoniae* or an isogenic ZomB mutant. At specific time points post colonisation the upper respiratory tract was lavaged with sterile PBS and the bacterial burdens in the lavage fluid quantified (**a**) and the lungs removed, homogenised and the bacterial burdens quantified to determine lower respiratory tract colonisation levels (**b**). The data from each mouse and sample are provided with the mean CFU +/− the standard error of the mean indicated. Statistical analysis was performed using a Kruskal-Wallis test with Dunn’s Multiple Comparisons. \**P* < 0.05.

## Discussion

Through the application of a functional genomics approach to a large collection of sequenced *S. pneumoniae* isolates, we have identified >100 novel putative affectors of Ply activity (supp. tables 1-3). Of these associated loci we have determined the molecular detail of the interaction between the ZomB protein encoded on ICE*Sp*23FST81 and Ply activity, where ZomB acts as a positive transcriptional regulator of the *ply* operon. We demonstrate that ZomB has ATP dependent nuclease activity and that it can bind with high affinity to multiple regions within the *ply* locus. Further work is required? to understand how the binding of ZomB to this locus affect its transcription, but we hypothesis its role is in the unravelling of secondary structures that would otherwise limit pneumolysin gene transcription.

Amongst the other loci identified by all three GWAS approaches as associated with Ply production were *bgaA* and *msmG*, two genes involved in carbohydrate utilisation. BgaA is a surface expressed b-galactosidase known to play a role in pneumococcal growth, resistance to opsonophagocytic killing, and adherence [48], whereas MsmG is part of the multiple sugar metabolism system [49]. A close relationship between metabolism and virulence is well established for other bacterial pathogens [50–52], and this has been recently established for the pneumococci where a clear link between capsule production and metabolism has been established. This work suggests that the effect of pneumococcal metabolism on its virulence may extend beyond capsule production and include an effect of Ply production, which is currently under investigation.

In this work we have identified and characterised a gene encoded on an MGE with specific and targeted activity for an operon that is critical to several aspects of the biology of *S. pneumoniae*. The PMEN1 lineage is believed to have emerged from a relatively unremarkable background lineage (CC66) to become a globally successful pathogenic lineage, and this is at least partially attributed to the acquisition of ICESp23FST81 [8,17,18]. This MGE, which is present across the whole PMEN1 lineage, confers resistance to tetracycline, macrolides and chloramphenicol and thus provides a clear benefit upon exposure to these antibiotics. However, for many bacterial species antibiotic resistance often incurs a fitness cost in the absence of the antibiotic, which can be offset, for example, by reducing the energetically costly production of toxins [28,29]. Here, in stark contrast, we observe the simultaneous acquisition of increased resistance with increased toxin production. We believe this work has uncovered an intriguing form of interdependency between a host bacterium and an MGE, where increased Ply expression due to the contribution this makes to transmission, has potentially converted a strain from being a stable coloniser to an efficient transmitter. The long-term benefit the contribution ZomB makes to this conversion may override the short-term increased energetic costs, potentially resulting in the evolution of this globally successful pneumococcal lineage.

## Authors Statements

### Authors contributions

EJS, DJM & DB develop methodology, performed experiment, analysed data and contributed to writing the manuscript. SD & TB provided support and supervision. MR analysed data and contributed to writing the manuscript. JAL, NJC, DJW, SJE & RD analysed data. SB & NJC provided resources. AN, HJ, TvO and DT provided guidance, expertise and resources. OJW and MSD performed experiments and provided advice on helicase proteins. SC and RMM designed performed and analysed the data from the animal experiments, and contributed to writing the manuscript. RCM conceptualised the project, developed the methodology, secured the funding, provided supervisory oversight and wrote the manuscript.

## Acknowledgements

We would also like to thank Prof. Tim Mitchell for helpful discussions during the preparation of this manuscript.

## REFERENCES

1. Deng, X. et al. Whole-genome sequencing reveals the origin and rapid evolution of an emerging outbreak strain of *Streptococcus pneumoniae* 12F. Clin Infect Dis 2016;62:1126–1132.

2. Ioachimescu, O.C., Ioachimescu, A.G. & Iannini, P.B. Severity scoring in community acquired pneumonia caused by *Streptococcus pneumoniae:* a 5-year experience. Int J Antimicrob Agents 2004;24:485–490.

3. O’Brien, K.L. et al. Burden of disease caused by *Streptococcus pneumoniae* in children younger than 5 years: global estimates. Lancet 2009;374:893–902.

4. Canvin, J.R. et al. The role of pneumolysin and autolysin in the pathology of pneumonia and septicaemia in mice infected with a type 2 pneumococcus. J Infect Dis 1995;172:119–123.

5. Hirst, R.A. et al. The role of pneumolysin in pneumococcal pneumonia and meningitis. Clin Exp Immunol 2004;138:195–201.

6. Hirst, R.A. et al. *Streptococcus pneumoniae* deficient in pneumolysin or autolysin has reduced virulence in meningitis. J of Infect Dis 2008;197;744–751.

7. Mitchell, A.M. & Mitchell, T.J. *Streptococcus pneumoniae:* virulence factors and variation. Clin Microbiol Infect Dis 2010;16:411–418.

8. Croucher, N.J. et al. Role of Conjugative Elements in the Evolution of the Multidrug-Resistant Pandemic Clone *Streptococcus pneumoniae* Spain23F ST81. J Bacteriol 2009;191:1480–9.

9. Mitchell, T.J. et al. Complement activation and antibody binding by pneumolysin via a region of the toxin homologous to a human acute-phase protein. Mol Microbiol 1991;5: 1883–8.

10. Paton, J.C., Rowan-Kelly, B. & Ferrante, A. Activation of human complement by the pneumococcal toxin pneumolysin. Infect Immun 1984;43:1085–7.

11. Subramanian, K. et al. Pneumolysin binds to the mannose receptor C type 1 (MRC-1) leading to anti-inflammatory responses and enhanced pneumococcal survival. Nat Microbiol 2019;4:62–70.

12. Kadlioglu, A., et al. Upper and lower respiratory tract infection by Streptococcus pneumoniae is affected by pneumolysin deficiency and differences in capsule type. Infect Immun 2002;70:2886–90.

13. van Rossum, A.M., Lysenko, E.S. & Weiser. J.N. Host and bacterial factors contributing to the clear-ance of colonization by *Streptococcus pneumoniae* in a murine model. Infect Immun 2005;73:7718–26.

14. Zafar, M.A., Wang, Y., Hamaguchi, S. & Weiser, J.N. Host-to-Host Transmission of Streptococcus pneumoniae Is Driven by Its Inflammatory Toxin, Pneumolysin. Cell Host Microbe 2017;21:73–83.

15. Riegler, A.N. Brissac, T., Gonzalez-Juarbe, N. & Orihuela C.J. Necroptotic Cell Death Promotes Adaptive Immunity Against Colonizing Pneumococci. Front Immunol 2019;10:615.

16. Anderson R, & Feldman C. Pneumolysin as a potential therapeutic target in severe pneumococcal disease. J Infect 2017;74:527–544.

17. Croucher, N.J. et al. Rapid pneumococcal evolution in response to clinical interventions. Science 2011;331:430–4.

18. Wyres, K.L. et al. The multidrug-resistant PMEN1 pneumococcus is a paradigm for genetic success. Genome Biology 2012;13:R103.

19. Chen, L., Ge, X. & Xu, P. Identifying essential *Streptococcus sanguinis* genes using genome-wide deletion mutation. Methods Mol. Biol 2015;1279:15–23.

20. Macrina, F.L., Tobian, J.A., Jones, K.R., Evans, R.P. & Clewell D.B. A cloning vector able to replicate in Escherichia coli and Streptococcus sanguis. Gene 1982;193:345–53.

21. Earle, S.G. et al. Identifying lineage effects when controlling for population structure improves power in bacterial association studies. Nat. Microbiol 2016;1:16041.

22. Lees, J.A. et al. pyseer: a comprehensive tool for microbial pangenome-wide association studies. Bioinformatics 2018;34:4310–2.

23. Livak, K.J. & Schmittgen, T.D. Analysis of relative gene expression data using real time quantitative PCR and the 2(-Delta Delta C(T)). Methods 2001;25:402–8.

24. Langmead, B. & Salzberg, S.L. Fast gapped-read alignment with Bowtie 2. Nat. Methods 2012;9:357–359.

25. Heng, L. et al. 1000 Genome Project Data Processing Subgroup, The Sequence Alignment/Map format and SAMtools. Bioinformatics 2009;25:2078–9.

26. Liao, Y., Smyth, G.K. & Shi W. FeatureCounts: an efficient general purpose program for assigning sequence reads to genomic features. Bioinformatics 2014;30:923–930.

27. Love, M.I., Huber, W. & Anders, S. Moderated estimation of fold change and dispersion for RNA-seq data with DESeq2. Genome Biol 2014;5:550.

28. Rudkin, J.K. et al. Methicillin resistance reduces the virulence of healthcare-associated methicillin-resistant *Staphylococcus aureus* by interfering with the *agr* quorum sensing system. J Infect Dis 2012;205:798–806.

29. Collins, J. et al. Offsetting virulence and antibiotic resistance costs by MRSA. ISME J 2010;4:577–84.

30. Hraiech, S. et al. Impaired virulence and fitness of a colistin-resistant clinical isolate of *Acinetobacter baumannii* in a rat model of pneumonia. Antimicrob. Agents Chemother 2013;57:5120–1.

31. Linares, J.F., et al. Overexpression of the multidrug efflux pumps MexCD-OprJ and MexEF-OprN is associated with a reduction of type III secretion in *Pseudomonas aeruginosa*. J. Bacteriol 2005;187:1384–91.

32. Azoulay-Dupuis, E. et al. Relationship between Capsular Type, Penicillin Susceptibility, and Virulence of Human *Streptococcus pneumoniae* Isolates in Mice. Antimicrob. Agents Chemother 2000;44:1575–7.

33. Rieux, V., Carbon, C. & Azoulay-Dupuis, E. Complex relationship Between Acquisition of Beta-Lactam Resistance and Loss of Virulence in *Streptococcus Pneumoniae*. J Infect Dis 2001;184:66–72.

34. Greene, N.G., Narciso, A.R., Filipe, S.R. & Camilli, A. Peptidoglycan Branched Stem Peptides Contribute to Streptococcus pneumoniae Virulence by Inhibiting Pneumolysin Release. PLoS Pathog 2015;11:e1004996.

35. Zhang, J., Kasciukovic, T. & White, M.G. The CRISPR Associated Protein Cas4 Is a 5’ to 3’ DNA Exonuclease with an Iron-Sulfur Cluster. PLoS One 2012;7:e47232.

36. Sampson, T.R. et al. A CRISPR/Cas system mediates bacterial innate immune evasion and virulence. Nature 2013;497;254–7.

37. Heidrich, N. et al. The CRISPR/Cas system in *Neisseria meningitidis* affects bacterial adhesion to human nasopharyngeal epithelial cells. RNA Biol 2019;16:390–396.

38. Ma, K. et al. cas9 Enhances Bacterial Virulence by Repressing the regR Transcriptional Regulator in *Streptococcus agalactiae*. Infect Immun 2018;86:e00552–17.

39. Gilhooly, N.S., Gwynn, E.J. & Dillingham, M.S. Superfamily 1 Helicases. Front Biosci 2013;5:206–16.

40. Fernebro, J. et al. The influence of in vitro fitness defects on pneumococcal ability to colonize and to cause invasive disease. BMC Microbiol 2008;8:65.

41. Kimaro Mlacha, S.Z., et al. Phenotypic, genomic, and transcriptional characterization of *Streptococcus pneumoniae* interacting with human pharyngeal cells. BMC Genomics 2013;14:383.

42. Croucher, N.J., Vernikos, G.S., Parkhill, J. & Bentley, S.D. Identification, variation and transcription of pneumococcal repeat sequences. BMC Genomics 2011;12:120.

43. Knutsen, E. et al. BOX Elements Modulate Gene Expression in *Streptococcus Pneumoniae:* Impact on the Fine-Tuning of Competence Development. J Bacteriol 2006;188:8307–12.

44. McCarville, J.L. & Ayres, J.S. Disease tolerance: concept and mechanisms. Current opinion in immunology 2018;50:88–93.

45. Neill, D.R. et al. Density and duration of pneumococcal carriage is maintained by transforming growth factor ß1 and T regulatory cells. Am J Respir Crit Care Med 2014;89:1250–9.

46. Wolf, A.I. et al. Pneumolysin Expression by Streptococcus pneumoniae Protects Colonized Mice from Influenza Virus-induced Disease. Virology 2014;0:254–265.

47. Inomataa, M. et al. Macrophage LC3-associated phagocytosis is an immune defense against Streptococcus pneumoniae that diminishes with host aging. PNAS 2020;117:33561–33569.

48. Singh AK., et al. Unravelling the multiple functions of the architecturally intricate Streptococcus pneumoniae beta-galactosidase, BgaA. PLoS Pathog 2014;0:e1004364.

49. Russell RR., et al. A binding protein-dependent transport system in Streptococcus mutans responsible for multiple sugar metabolism. J Biol Chem 1992;267:4631–7.

50. Stevens E., Laabei M, Gardner S, Sommerville GA, Massey RC. Cytolytic toxin production by Staphylococcus aureus is dependent upon the activity of the protoheme IX farnesyltransferase. Sci Rep 2017;7:13744.

51. Bouillaut L, Dubois T, Sonenshein AL, Dupuy B. Integration of metabolism and virulence in Clostridium difficile. Res Microbiol 2015;166:375–83.

52. Le Bouguénec C, Schouler C. Sugar metabolism, an additional virulence factor in enterobacteria. Int J Med Microbiol 2011;301:1–6.

